# GRASP: Graph Representation Learning with Assay Supervision for Molecular Properties

**DOI:** 10.64898/2026.09.28.755019

**Authors:** Satya Pratik Srivastava, Rohan Gorantla, Sharath Krishna Chundru, Harshit Singh, Antonia S. J. S. Mey, Rajeev Kumar Singh

## Abstract

Molecular structures are abundant, but measured properties are scarce and scattered across assays that each cover different compounds. Most molecular pretraining pipelines use only the structures, moving directly from self-supervision to endpoint-specific fine-tuning and leaving large, sparsely labelled bioactivity databases unused. We present GRASP, a 93.5M-parameter graph Transformer trained in two stages: replaced-token detection over atom tokens with shortest-path relative attention using 1.54B molecule presentations from ZINC20, followed by adaptation on sparse labels from 642 ChEMBL assays. On 23 cluster-held-out OpenADMET endpoints, fine-tuned GRASP reaches a mean MAE of 0.374, compared with 0.383 for CheMeleon and 0.408 for ECFP4-LightGBM, and has lower error than Chemprop on 22 of 23 endpoints. Introducing the intermediate ChEMBL stage reduces mean MAE from 0.407 to 0.374 and improves 21 of 23 endpoints. In a controlled smaller-scale recipe comparison, replaced-token detection yields lower MAE than a masked-token recipe on 16 of 23 OpenADMET endpoints and favorable point estimates on 17 of 22 TDC ADMET tasks. Sparse multi-assay data, used as an intermediate training stage, is an effective complement to structural pretraining.

## Introduction

Learning useful molecular representations is difficult because we have far more data about molecular structures than about experimentally measured properties [1, 2]. Databases such as ZINC contain enormous numbers of molecular structures [3], but experimental property data is available for only a small fraction of these molecules. Obtaining it is expensive because each measurement requires laboratory experiments, so the data remains sparse. It is also spread across many different assays, each covering a different set of molecules, with little overlap between them [4]. Most molecular pretraining pipelines use the first resource for self-supervision and then move directly to endpoint-specific fine-tuning. This leaves a valuable source of supervision underutilised: large bioactivity databases such as ChEMBL, whose incomplete assay matrices still connect molecular structure to experimentally observed chemical behaviour [5, 6].

We introduce Graph Representation Learning with Assay Supervision for Molecular Properties - GRASP, a 93.5M-parameter graph Transformer that learns from these sources in sequence. The first stage learns molecular structure through replaced-token detection (RTD): a compact generator proposes contextual atom-token replacements, and a discriminator predicts which atoms were replaced [7]. Each atom is represented by a radius-0 Morgan token, while shortest-path relative attention supplies molecular topology [8–10]. Structural training uses 1.54B molecular presentations from ZINC20. The second stage adapts the encoder using approximately 512,000 molecules and 642 sparsely observed ChEMBL assays. The resulting model can then be adapted to predict a specific molecular property by full fine-tuning, LoRA, or frozen layer mixing [11].

On 23 cluster-held-out OpenADMET endpoints [12–14], full fine-tuning reaches a mean MAE of 0.374, compared with 0.383 for CheMeleon, 0.408 for tuned ECFP4-LightGBM, and 0.440 for Chemprop D-MPNN. GRASP has lower endpoint MAE than CheMeleon on 12 of 23 endpoints, ECFP4-LightGBM on 16 of 23, and Chemprop on 22 of 23. With frozen encoders, GRASP also improves on Mol-JEPA on 18 of 23 endpoints under the same splits and regression protocol.

In the matched stage comparison, ChEMBL adaptation lowers mean MAE from 0.407 to 0.374 and improves 21 endpoints, in GRASP-Small, RTD improves on the implemented masked-token objective on 16 endpoints. We further evaluate the model on the 22-task TDC ADMET benchmark, which covers both classification and regression tasks [15]. ChEMBL adaptation yields favorable point estimates on 19 of 22 tasks (Section ; Table A13). Overall, the results show that learning molecular structure at scale and then adapting on sparse experimental data improves performance on downstream property prediction.

## Related Work

Controlled studies show that molecular pretraining gains depend on features, architecture, data splits, and downstream optimization [16], while scaling studies report benefits from larger and more diverse supervision [17, 18]. Molecular representation learning has consequently explored diverse signals, including self-supervised graph objectives (GROVER) [19], contrastive learning (MolCLR) [20], molecular strings (ChemBERTa, MoLFormer) [21, 22], geometry (Uni-Mol/Uni-Mol2) [23, 24], descriptors (CheMeleon) [25], and bioactivity supervision (MolE) [26]. GRASP instead combines shortest-path relative attention [9, 10] with ELECTRA-style replaced-token detection and gradient-disentangled embedding sharing [7, 27]; replacement-based graph objectives have also been studied in DiscoGNN [28]. Rather than attributing performance to individual architectural components, we test two controlled questions: whether RTD improves over masked-token pretraining, and how sparse multi-assay bioactivity adaptation changes the resulting representation.

## Overview of GRASP

### Molecular representation and encoder

We tokenize each molecule with RDKit radius-0 Morgan hashes, yielding one atom token from a 211-token vocabulary including PAD, MASK, UNK, and CLS [8, 29]. The input also contains a relative-distance index *r*(*i, j*) derived from shortest-path bond distance. For connected atom pairs, *r*(*i, j*) = *δ*(*i, j*)+1, where *δ*(*i, j*) is their shortest-path bond distance. We set *r*(*i, j*) = 0 for pairs involving CLS or padding and for atom pairs in disconnected components. The discriminator is a 12-layer graph Transformer with hidden size 768, 12 heads, feed-forward size 3,072, and 93,484,800 encoder parameters [30]. It uses content-to-position relative attention,

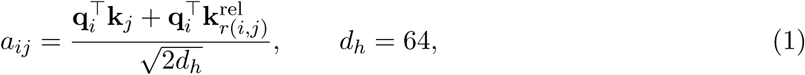

together with learned absolute positional embeddings based on the RDKit atom ordering, while the attention scores themselves remain content-based.

### Graph replaced-token detection

Figure 1 summarizes the objective.

**Figure 1.**
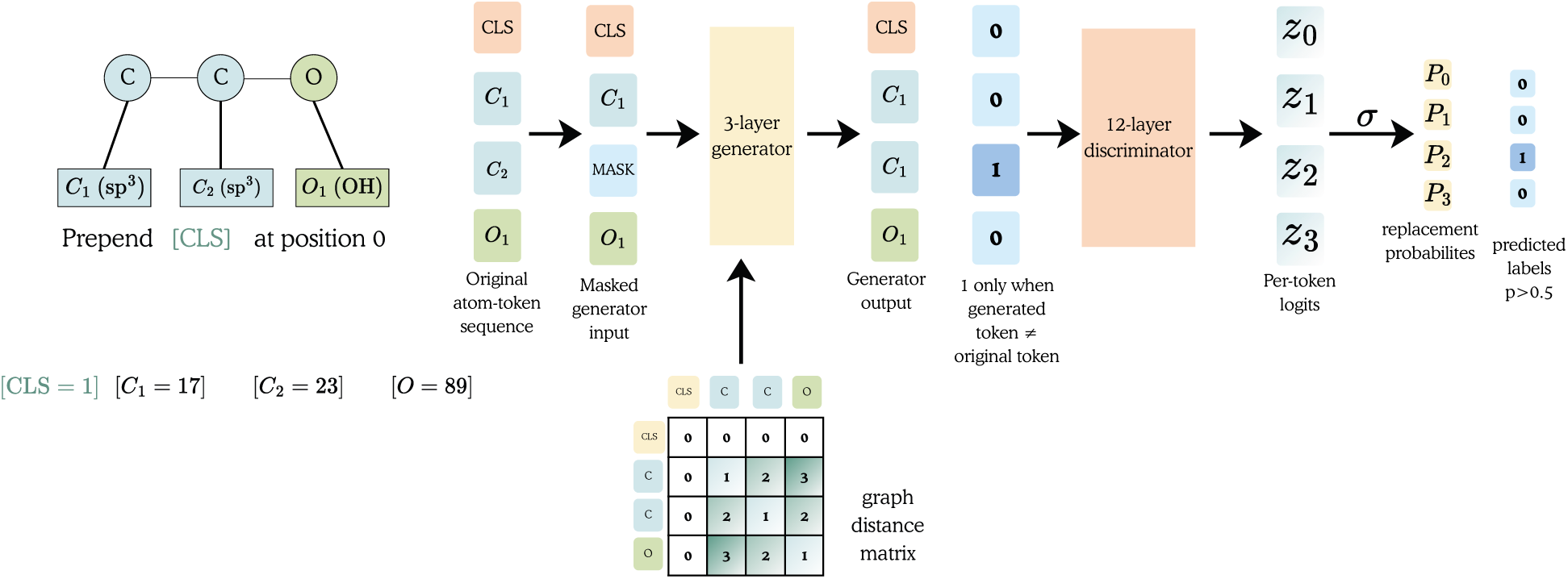
Graph replaced-token detection. Each heavy atom is tokenized as a radius-0 Morgan hash and a CLS token is prepended at position 0. Atom tokens are independently selected with probability 0.25 and replaced by MASK, and a three-layer generator predicts the original token at each selected position. The generator’s argmax prediction is written back into the graph, so a position is labelled as replaced only when the generated token differs from the original. The 12-layer discriminator receives the resulting token sequence together with the shortest-path graph-distance matrix and predicts, at every valid position, whether that token was replaced.

Each atom token is independently selected with probability 0.25 and replaced by MASK. A three-layer, 256-hidden generator predicts the original token, and its argmax prediction is inserted into the graph. For selected positions *M*, generator cross-entropy is

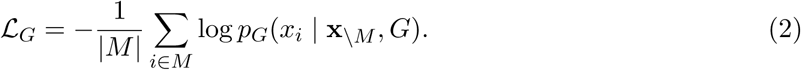

The discriminator predicts at every valid position *T*, including CLS, whether the realized input differs from the original: *r_i_* = ⊮[*x̃_i_* ≠ *x_i_*], with *r*_CLS_ = 0. Its loss is

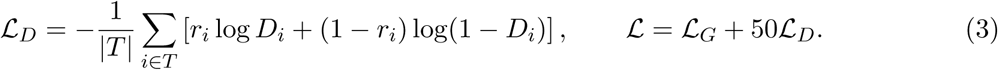

Gradient-disentangled embedding sharing implements *W_D_* = stopgrad(*W_G_*) + *B_D_*: the generator supplies shared token-embedding values, while discriminator gradients update only the learned bias *B_D_*.

### Two-stage training

Step 1 applies RTD to 1,538,240,768 molecular presentations from 1,538,240,669 ZINC20 records over 6,133,753 optimizer updates [3]. Step 2 fine-tunes the encoder for 80,000 updates on 511,898 ChEMBL 36 molecules and 642 binary binding or functional assays [5, 6]. Labels come from activity comments or pChEMBL thresholds (≥5.5 active, ≤4.5 inactive); contradictory pairs are removed and assays require at least 500 molecules. A linear 642-output head reads the final-layer CLS state, and binary cross-entropy is computed only for observed labels. The fixed 50,000-step encoder snapshot is used downstream.

### Downstream adaptation

For OpenADMET, the three adaptation modes use a softmax-weighted mixture of layers 4, 8, 10, and 12, additive attention pooling over atom tokens (excluding CLS), and a 512-hidden MLP head. Full fine-tuning updates the entire encoder; frozen mixing keeps the encoder fixed; LoRA [31] keeps the base weights fixed and adds rank-16 query/value updates (*α* = 32, dropout 0.1) in every attention layer. The secondary TDC evaluation uses a separate task-specific protocol based on the final-layer CLS representation, described in Appendix A.2.

## Results

### GRASP improves molecular property prediction across diverse ADMET endpoints

#### Task

We evaluate molecular property prediction on OpenADMET, a multi-endpoint benchmark spanning clearance, solubility, permeability, plasma and tissue binding, efflux, PXR activity, and antiviral Mpro potency (pIC50 or pEC50). Given a molecular graph, the model predicts the continuous value of each endpoint independently. Performance is measured using mean absolute error (MAE), with lower values indicating better predictive performance.

### Dataset and Evaluation

OpenADMET comprises 23 endpoints, with 43,881 labelled measurements across 14,261 unique SMILES. Each endpoint is evaluated on three reconstructed cluster-held-out splits, and we report the test MAE averaged across the three splits. Dataset sizes, target transformations, and endpoint-specific details are provided in Appendix A.1 Table A4.

### Baselines

We compare full fine-tuning (FT) with ECFP4-LightGBM and RDKit-LightGBM [**?**], ECFP4 and RDKit random forests, Chemprop [**?**], and CheMeleon, all evaluated on identical partitions. For the frozen-encoder comparison, we use each model’s architecture-native molecular representation: Mol-JEPA’s released CLS embedding [**?**] and GRASP’s learned mixture of layers 4, 8, 10, and 12 followed by atom-attention pooling. Both representations are evaluated using the same regression MLP and the same data partitions and downstream training protocol. Thus, the downstream evaluation is matched, while the representation readout remains architecture-specific.

### Performance

GRASP full fine-tuning reaches a mean MAE of 0.374 ±0.003 across the 23 endpoints, the lowest of the methods compared (Table 1). It improves on Chemprop D-MPNN by 14.9% (22 of 23 endpoints) and on tuned ECFP4-LightGBM by 8.2% (16 of 23). The margin over CheMeleon, the strongest baseline, is smaller: 2.4% in the mean, with lower MAE on 12 of 23 endpoints. Per-endpoint results for all methods are given in Table A2 and paired contrasts in Table A3; Figure 2 shows the corresponding per-endpoint ranks and two representative fits. GRASP FT also had the lowest mean MAE among adaptation strategies (FT: 0.374; LoRA: 0.380; frozen mixing: 0.387), with evidence of a difference only against frozen mixing after correction (*p* = 0.037). Under this common downstream evaluation protocol, GRASP also improved over Mol-JEPA (0.387 vs. 0.412; 18/23 endpoints; ΔMAE = −0.025, 95% CI [−0.042, −0.008]).

**Figure 2.**
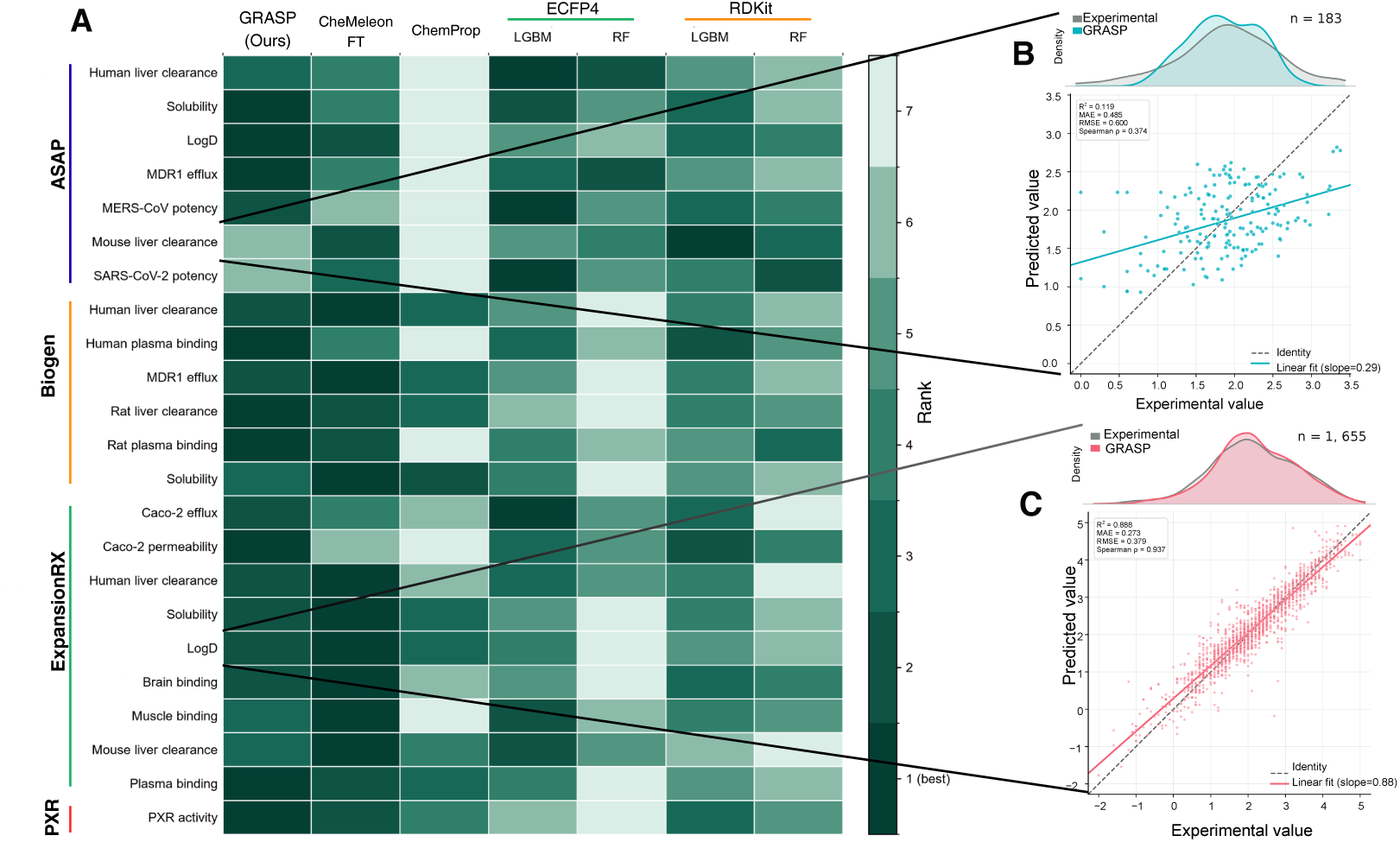
OpenADMET-23 performance. **(A)** Rank of each method on each of the 23 cluster-held-out endpoints, from 1 (best, dark) to 7, with endpoints grouped by source. Ranks are computed from the endpoint mean MAE over the three splits reported in Table A2. **(B, C)** Predicted against experimental values for GRASP full fine-tuning on two endpoints, pooled over the three splits, illustrating the range of difficulty in the panel: **(B)** ASAP mouse liver clearance (*n* = 183), among the weakest endpoints for every method, and **(C)** ExpansionRx LogD (*n* = 1,655), among the strongest. Marginal densities compare the experimental and predicted distributions. The dashed line is identity and the solid line is the least-squares fit; pooled MAE differs slightly from the split-averaged MAE in Table A2 because splits have unequal test sizes.

**Table 1.** OpenADMET-23 summary. % impr. is the relative reduction in mean MAE achieved by GRASP full fine-tuning (FT) over the comparator, (1 − MAE_FT_*/*MAE_comp_) ×100, and Wins/23 counts endpoints where FT has lower MAE. In the frozen-encoder block, the comparison is instead GRASP frozen mixing versus Mol-JEPA. All comparisons use identical reconstructed cluster-held-out splits.

| Comparator | Mean MAE | % impr. | Wins/23 |
| --- | --- | --- | --- |
| *Rerun baselines (vs. FT, $0.374 \pm 0.003$)* |  |  |  |
| ECFP4-LightGBM | $0.408 \pm 0.014$ | 8.2 | 16 |
| RDKit-LightGBM | $0.412 \pm 0.011$ | 9.2 | 21 |
| RDKit random forest | $0.431 \pm 0.001$ | 13.1 | 21 |
| Chemprop D-MPNN | $0.440 \pm 0.016$ | 14.9 | 22 |
| ECFP4 random forest | $0.453 \pm 0.001$ | 17.3 | 20 |
| CheMeleon FT | $0.383 \pm 0.008$ | 2.4 | 12 |
| *Adaptation of GRASP (vs. FT)* |  |  |  |
| LoRA | 0.380 | 1.6 | 14 |
| Frozen layer mixing | 0.387 | 3.4 | 18 |
| *Frozen encoder (vs. GRASP frozen, 0.3869)* |  |  |  |
| Mol-JEPA (CLS + MLP) | 0.4123 | 6.2 | 18 |

**Table 2.** OpenADMET adaptation comparison. Mean MAE averages the 23 endpoint means across three cluster-held-out splits; ±denotes population standard deviation across split-wise macro-averages where available. Pairwise *p*-values compare each method with full fine-tuning and are Holm-adjusted across all three adaptation contrasts. The LoRA–frozen contrast is included in the correction but is not shown. The overall Friedman test gives *χ*^2^(2) = 7.91, *p* = 0.019.

| Method | Mean MAE ↓ | $p_{\text{Holm}}$ vs. FT |
| --- | --- | --- |
| **Full fine-tuning** | **$0.374 \pm 0.003$** | — |
| LoRA | 0.380 | 0.268 |
| Frozen layer mixing | 0.387 | 0.037 |

### RTD provides gains over MLM for pretraining

#### Task

We compare replaced-token detection (RTD) with masked language modeling (MLM) for GRASP pretraining. Both models use the same encoder architecture, pretraining data, ChEMBL adaptation stage, and downstream evaluation. The models differ only in the pretraining objective. RTD detects replaced radius-0 atom tokens. MLM predicts masked radius-2 atom-environment tokens.

### Dataset and Evaluation

Training a second model at the full 1.538B-presentation scale requires substantial compute. We therefore compare RTD and MLM with the smaller GRASP-Small configuration. Both models use 415M ZINC molecule presentations and the same 12-layer encoder with a hidden size of 768. The encoder has approximately 89M parameters. Both models use a 15% selected-token rate. Both models then use the same ChEMBL adaptation stage with 641 assays and 80,000 updates.

We freeze both encoders for the downstream evaluation. We evaluate them on 23 OpenADMET endpoints with the same three data splits. Both models use the same layer-mixing readout and regression head. Table 3 reports the test MAE averaged across the three splits.

**Table 3.** Frozen-transfer comparison of RTD and MLM recipes on OpenADMET-23. RTD-15%-S2 and MLM-15%-S2 both use GRASP-Small with a 15% selected-token rate. Both use radius-0 input tokens; MLM predicts radius-2 environments. After ChEMBL adaptation, both encoders are frozen and evaluated with the same learned layer-mixing readout and regression head. Values are mean MAE ± population standard deviation across three split-specific fits. Bold marks the lower unrounded mean. Δ = 100(MAE_MLM_ − MAE_RTD_)*/*MAE_MLM_; positive values favor RTD.

| Endpoint | GRASP-RTD | GRASP-MLM | $\Delta$ (%) |
| --- | --- | --- | --- |
| **ASAP** |  |  |  |
| Human liver clearance | 0.38 ± .04 | **0.34 ± .10** | −10.81 |
| Solubility | **0.39 ± .04** | 0.42 ± .13 | +6.36 |
| LogD | **0.59 ± .06** | 0.67 ± .15 | +11.59 |
| MDR1 efflux | **0.37 ± .02** | 0.39 ± .02 | +5.30 |
| MERS-CoV potency | **0.60 ± .08** | 0.60 ± .12 | +0.66 |
| Mouse liver clearance | 0.44 ± .04 | **0.43 ± .06** | −2.68 |
| SARS-CoV-2 potency | **0.64 ± .09** | 0.71 ± .12 | +10.32 |
| **Biogen** |  |  |  |
| Human liver clearance | **0.37 ± .03** | 0.38 ± .00 | +4.24 |
| Human plasma binding | 0.46 ± .16 | **0.45 ± .18** | −1.79 |
| MDR1 efflux | **0.42 ± .01** | 0.43 ± .01 | +2.91 |
| Rat liver clearance | **0.42 ± .02** | 0.43 ± .07 | +2.19 |
| Rat plasma binding | 0.54 ± .07 | **0.49 ± .05** | −9.84 |
| Solubility | **0.35 ± .03** | 0.42 ± .05 | +16.50 |
| **ExpansionRx** |  |  |  |
| Caco-2 efflux | **0.24 ± .01** | 0.26 ± .01 | +6.06 |
| Caco-2 permeability | **0.35 ± .04** | 0.35 ± .05 | +1.45 |
| Human liver clearance | **0.39 ± .02** | 0.40 ± .01 | +1.16 |
| Solubility | **0.51 ± .05** | 0.51 ± .04 | +1.34 |
| LogD | **0.42 ± .04** | 0.42 ± .02 | +0.96 |
| Brain binding | **0.28 ± .01** | 0.30 ± .03 | +6.93 |
| Muscle binding | 0.25 ± .04 | **0.25 ± .04** | −0.31 |
| Mouse liver clearance | 0.38 ± .02 | **0.38 ± .03** | −0.10 |
| Plasma binding | 0.32 ± .03 | **0.30 ± .05** | −5.63 |
| **PXR** |  |  |  |
| PXR activity | **0.55 ± .04** | 0.58 ± .03 | +6.48 |
| **Mean** | **0.42 ± .01** | 0.43 ± .01 | +2.83 |

### Performance

RTD achieves lower mean MAE than GRASP-MLM (0.418 vs. 0.431), outperforming it on 16/23 endpoints as presented in Table 3. The paired mean improvement is 0.0122 MAE (Holm-adjusted exact Wilcoxon *p* = 0.0969). GRASP-RTD wins all five LogD and solubility endpoints, with the largest gains on Biogen solubility (+16.5%), ASAP LogD (+11.6%), and SARS-CoV-2 potency (+10.3%). MLM’s seven wins are all clearance or binding endpoints, largest on ASAP human liver clearance (−10.8%) and Biogen rat plasma binding (−9.9%).

### ChEMBL bioactivity adaptation improves molecular property prediction

#### Task

We test the effect of the ChEMBL bioactivity adaptation stage. GRASP-S1 is the Step-1 checkpoint trained with structural RTD only. GRASP also completes Step-2 adaptation on 642 sparsely observed ChEMBL assays. Step 2 includes both additional training on ChEMBL molecules and supervision from measured assay values. Therefore, this comparison measures the effect of the complete adaptation stage rather than assay supervision alone.

### Dataset and Evaluation

We fully fine-tune both models on the same 23 OpenADMET endpoints. We evaluate each endpoint on the same three cluster-held-out splits. We report the test MAE averaged across the three splits.

### Performance

ChEMBL adaptation lowers the mean MAE from 0.4069 to 0.3742, an 8.0% reduction. GRASP gives a lower MAE on 21 of 23 endpoints Table 4). The largest relative gains occur for ASAP solubility (24.2%), ExpansionRx plasma binding (19.5%), ASAP LogD (16.4%), and ExpansionRx brain binding (14.3%). GRASP-S1 gives a lower MAE on two endpoints: ASAP mouse liver clearance (6.7%) and ExpansionRx muscle binding (1.0%).

**Table 4.** Effect of ChEMBL bioactivity adaptation on OpenADMET-23. GRASP-S1 uses structural pretraining; GRASP additionally includes ChEMBL adaptation. Both checkpoints are fully fine-tuned on the same three cluster-held-out splits. Entries report mean test MAE ± population standard deviation across split-specific fits. The mean-row deviation is calculated from the three split-wise macro-average MAEs. Bold marks the lower unrounded mean. Δ = 100(MAE_S1_ − MAE_GRASP_)*/*MAE_S1_; positive values favor GRASP.

| Endpoint | GRASP-S1 | GRASP | $\Delta$ (%) |
| --- | --- | --- | --- |
| **ASAP** |  |  |  |
| Human liver clearance | 0.38 ± .07 | **0.36 ± .07** | +7.66 |
| Solubility | 0.42 ± .08 | **0.32 ± .05** | +24.21 |
| LogD | 0.56 ± .14 | **0.47 ± .06** | +16.35 |
| MDR1 efflux | 0.35 ± .03 | **0.32 ± .03** | +6.35 |
| MERS-CoV potency | 0.58 ± .11 | **0.56 ± .11** | +4.43 |
| Mouse liver clearance | **0.46 ± .04** | 0.49 ± .05 | -6.71 |
| SARS-CoV-2 potency | 0.66 ± .03 | **0.62 ± .05** | +6.50 |
| **Biogen** |  |  |  |
| Human liver clearance | 0.37 ± .03 | **0.33 ± .01** | +12.34 |
| Human plasma binding | 0.52 ± .11 | **0.47 ± .13** | +9.25 |
| MDR1 efflux | 0.37 ± .03 | **0.34 ± .02** | +9.56 |
| Rat liver clearance | 0.41 ± .02 | **0.38 ± .02** | +8.54 |
| Rat plasma binding | 0.47 ± .03 | **0.45 ± .06** | +4.65 |
| Solubility | 0.38 ± .03 | **0.37 ± .00** | +2.67 |
| **ExpansionRx** |  |  |  |
| Caco-2 efflux | 0.25 ± .02 | **0.23 ± .01** | +7.68 |
| Caco-2 permeability | 0.32 ± .02 | **0.29 ± .02** | +8.14 |
| Human liver clearance | 0.38 ± .02 | **0.35 ± .02** | +6.93 |
| Solubility | 0.44 ± .03 | **0.42 ± .02** | +5.13 |
| LogD | 0.30 ± .02 | **0.27 ± .02** | +9.08 |
| Brain binding | 0.29 ± .01 | **0.25 ± .01** | +14.32 |
| Muscle binding | **0.26 ± .04** | 0.26 ± .01 | -0.97 |
| Mouse liver clearance | 0.37 ± .01 | **0.34 ± .03** | +7.44 |
| Plasma binding | 0.30 ± .02 | **0.24 ± .03** | +19.53 |
| **PXR** |  |  |  |
| PXR activity | 0.52 ± .03 | **0.49 ± .03** | +5.03 |
| **Mean** | 0.41 ± .01 | **0.37 ± .00** | **+8.03** |

### Secondary evaluations: TDC

Dataset composition, scaffold splits, hyperparameter selection, and evaluation metrics for the 22-task TDC ADMET panel are detailed in Appendix A.2.

### GRASP remains competitive across TDC ADMET tasks

#### Task

We evaluate GRASP on 22 TDC ADMET tasks (13 classification, 9 regression) covering absorption, distribution, metabolism, excretion, and toxicity.

### Dataset and Evaluation

We use the fixed test sets of the TDC ADMET benchmark group without applying an additional heavy-atom filter or molecular truncation. We select hyperparameters with five independently seeded scaffold splits of the benchmark training and validation data. Test labels remain unused during hyperparameter selection. After hyperparameter selection, we fully fine-tune GRASP in three independent training runs.

### Baseline

We compare with the released CheMeleon D-MPNN results.

### Performance

GRASP has the better point estimate on 13 and CheMeleon on 9 (Table 5). GRASP’s largest margins are half-life (0.455 vs. 0.325 Spearman), VDss (0.665 vs. 0.546), and bioavailability (0.687 vs. 0.579 AUROC). CheMeleon’s clearest lead is hERG (0.867 vs. 0.773 AUROC).

**Table 5.** GRASP on the 22-task TDC ADMET benchmark. GRASP values are mean ± population standard deviation over three seeds after validation-only hyperparameter selection.

| Task | Metric | GRASP | CheMeleon |
| --- | --- | --- | --- |
| **Absorption** |  |  |  |
| Caco-2 Wang | MAE ↓ | $0.39 \pm .01$ | **$0.38 \pm .03$** |
| Bioavailability Ma | AUROC ↑ | **$0.69 \pm .02$** | $0.58 \pm .02$ |
| Lipophilicity AstraZeneca | MAE ↓ | **$0.41 \pm .00$** | $0.46 \pm .01$ |
| Solubility AqSolDB | MAE ↓ | **$0.74 \pm .01$** | $0.89 \pm .11$ |
| HIA Hou | AUROC ↑ | $0.95 \pm .02$ | **$0.96 \pm .01$** |
| P-glycoprotein Broccatelli | AUROC ↑ | $0.92 \pm .01$ | **$0.93 \pm .01$** |
| **Distribution** |  |  |  |
| BBB Martins | AUROC ↑ | $0.88 \pm .02$ | **$0.89 \pm .02$** |
| PPBR AZ | MAE ↓ | **$7.30 \pm .35$** | $7.82 \pm .15$ |
| VDss Lombardo | Spearman ↑ | **$0.67 \pm .02$** | $0.55 \pm .01$ |
| **Metabolism** |  |  |  |
| CYP2C9 inhibition | AUPRC ↑ | **$0.79 \pm .00$** | $0.78 \pm .01$ |
| CYP2D6 inhibition | AUPRC ↑ | **$0.72 \pm .02$** | $0.68 \pm .00$ |
| CYP3A4 inhibition | AUPRC ↑ | **$0.89 \pm .01$** | $0.87 \pm .01$ |
| CYP2C9 substrate | AUPRC ↑ | $0.41 \pm .03$ | **$0.43 \pm .01$** |
| CYP2D6 substrate | AUPRC ↑ | $0.64 \pm .03$ | **$0.67 \pm .04$** |
| CYP3A4 substrate | AUROC ↑ | **$0.67 \pm .04$** | $0.61 \pm .01$ |
| **Excretion** |  |  |  |
| Half-life Obach | Spearman ↑ | **$0.46 \pm .06$** | $0.33 \pm .03$ |
| Clearance hepatocyte AZ | Spearman ↑ | **$0.47 \pm .02$** | $0.38 \pm .05$ |
| Clearance microsome AZ | Spearman ↑ | $0.60 \pm .02$ | **$0.60 \pm .02$** |
| **Toxicity** |  |  |  |
| LD50 Zhu | MAE ↓ | **$0.49 \pm .00$** | $0.53 \pm .01$ |
| hERG | AUROC ↑ | $0.77 \pm .03$ | **$0.87 \pm .01$** |
| Ames | AUROC ↑ | $0.83 \pm .01$ | **$0.85 \pm .01$** |
| DILI | AUROC ↑ | **$0.93 \pm .02$** | $0.90 \pm .03$ |
| **Favorable tasks** |  | **13/22** | 9/22 |

### RTD improves performance across TDC tasks

### Task and Evaluation

We compare RTD with MLM on TDC in the GRASP-Small configuration of section (415M ZINC molecules, 15% selected-token rate, 80,000-update ChEMBL stage). RTD detects replaced radius-0 tokens, whereas MLM predicts masked radius-2 environments, so this compares implemented recipes rather than a single loss. Scaffold partitions, downstream heads, training settings, and three seeds are identical.

### Performance

RTD has the better point estimate on 17 of 22 tasks and MLM on 5 (exact two-sided sign test, raw *p* = 0.0169, Holm-adjusted *p* = 0.0338;Table A12). RTD’s gains include microsomal clearance (0.589 vs. 0.467 Spearman), hERG (0.818 vs. 0.753 AUROC), and lipophilicity (0.459 vs. 0.505 MAE). MLM’s five wins are the three CYP substrate tasks, DILI, and P-glycoprotein. Complete task-level results are reported in Table A12.

### ChEMBL bioactivity adaptation improves transfer across TDC tasks

#### Task

We test the effect of the ChEMBL bioactivity adaptation stage on downstream transfer. GRASP-S1 contains only the structural RTD stage. GRASP also includes the ChEMBL adaptation stage with sparse assay supervision. This comparison measures the effect of the complete adaptation stage rather than assay supervision alone.

### Dataset and Evaluation

We evaluate both checkpoints on the same 22 TDC ADMET tasks. Both models use the same fixed test sets and downstream heads. We select hyperparameters with the same five scaffold splits of the training and validation data. Both models use the same three random seeds for downstream training.

### Performance

GRASP has the favorable point estimate over GRASP-S1 on 19 of the 22 tasks (Table A13). The table reports both checkpoints for every task, using the same GRASP values as Table 5. Representative gains include CYP3A4 substrate prediction (AUROC: 0.565 ±0.046 to **0**.**674** ±**0**.**035**), CYP2D6 inhibition (AUPRC: 0.624 ±0.009 to **0**.**716** ±**0**.**017**), hepatocyte clearance (Spearman: 0.408 ±0.042 to **0**.**471** ±**0**.**018**), and microsomal clearance (Spearman: 0.556 ±0.048 to **0**.**602** ±**0**.**017**). GRASP also reduces MAE on Caco-2 permeability (0.470 ±0.040 to **0**.**389** ±**0**.**009**), solubility (0.794 ±0.003 to **0**.**736 ± 0**.**009**), lipophilicity (0.449 ±0.011 to **0**.**406** ±**0**.**003**), and LD50 (0.576 ±0.002 to **0**.**486** ±**0**.**002**). GRASP-S1 retains the favorable point estimate on half-life, hERG, and HIA. Overall, the adapted GRASP checkpoint is favorable on most TDC endpoints, with gains spanning classification and regression tasks.

## Discussion, Conclusion and Limitations

GRASP demonstrates that a molecular encoder can benefit from learning structure at scale and then incorporating sparse experimental bioactivity. On OpenADMET-23, full fine-tuning reaches a mean MAE of 0.3742, compared with 0.4075 for ECFP4-LightGBM and 0.4395 for Chemprop. The endpoint counts are important: GRASP improves on ECFP4-LightGBM on 16 of 23 tasks (Holm-adjusted sign test *p* = 0.093; the paired Wilcoxon test over the same endpoints gives *p* = 0.006) and on Chemprop on 22 of 23. The result is therefore not only a lower macro-average; the model is useful across most of the evaluated clearance, solubility, permeability, binding, efflux, and activity endpoints. GRASP is the single best method on 9 of the 23 endpoints (Table A2), tuned fingerprint and descriptor models remain competitive per endpoint at these training-set sizes, and the advantage we report is a consistent aggregate one. On the separate TDC benchmark, GRASP has the favorable point estimate on 13 of 22 tasks with a comparable CheMeleon value, extending the evaluation to both classification and regression under scaffold holdout.

The frozen-encoder experiment shows that this performance is not limited to updating the full backbone. Using the same endpoint partitions and regression MLP, frozen GRASP obtains a mean MAE of 0.3869, compared with 0.4123 for Mol-JEPA, and improves on 18 of 23 endpoints. Downstream adaptation provides an additional gain: full fine-tuning reaches 0.3742, LoRA reaches 0.3800, and frozen layer mixing reaches 0.3869. LoRA updates only 1.04% of the parameters, making it a strong option when many endpoint-specific models must share one pretrained encoder. The difference between full fine-tuning and LoRA is not resolved after multiplicity correction, whereas full fine-tuning is separated from frozen mixing.

The controlled training comparisons support both stages of the recipe. In GRASP-Small, RTD improves on the implemented MLM objective on 16 of 23 OpenADMET endpoints and 17 of 22 TDC tasks. Adding sparse ChEMBL bioactivity adaptation lowers OpenADMET mean MAE from 0.4069 to 0.3742 and improves 21 of 23 endpoints relative to the Step-1 checkpoint. Individual endpoints do not always favor GRASP, but the main pattern is consistent across the primary comparison, the frozen-encoder analysis, the stage comparison, and the secondary benchmark. GRASP therefore provides a practical foundation for molecular property prediction: one graph encoder, trained from abundant structures and sparse assays, that supports both accuracy-oriented full fine-tuning and parameter-efficient adaptation.

This ability to adapt from limited experimental supervision could also support prospective molecular discovery, including low-data assay modelling and active-learning [32] or lab-in-the-loop workflows, where the encoder is continually updated as new measurements become available to prioritize subsequent compounds for evaluation.

### Limitations

Several conclusions are intentionally narrower than the full training recipe. First, the GRASP-S1–GRASP comparison evaluates the complete ChEMBL adaptation stage and does not isolate assay supervision from additional optimization or exposure to the ChEMBL molecular distribution. Second, the structural-overlap analysis tests molecular exposure but does not establish the absence of semantically related assay supervision between ChEMBL and downstream endpoints. Third, the RTD–MLM comparison is performed in GRASP-Small and compares implemented pretraining recipes rather than isolating pretraining scale. Fourth, the principal downstream experiments use the 50,000-step Step-2 snapshot without a systematic downstream checkpoint-sensitivity study. Finally, the TDC comparison with released CheMeleon scores is descriptive because the two methods were not trained and selected within a single paired experimental pipeline.

## Acknowledgement

The authors acknowledge Department of Computer Science and Engineering, Shiv Nadar University, Delhi-NCR and the Edinburgh International Data Facility (EIDF) for providing the computational and data storage resources used in this research

## AI Use Disclosure

Generative AI tools were used to assist with manuscript writing and editing. All AI-assisted content was reviewed and verified by the authors, who take full responsibility for the final work.

## Code and Model Availability

The implementation and model-release information are available at https://github.com/caithmac/GRASP.

## A Additional Experimental Details

### A.1 Architecture and Training Details

This appendix specifies the input representation, architecture, and training settings needed to reproduce the model.

RDKit uses radius 0, useFeatures=False, redundant environments enabled, and chirality disabled. The 32-bit hashes form 207 observed atom types plus PAD, MASK, UNK, and CLS (211 entries); unseen hashes map to UNK. These hashes are not reversible feature tuples. For connected atom pairs, graph distance *d* is stored as the relative-distance index *d* + 1; index 0 is reserved for pairs involving CLS or padding and for atom pairs in disconnected components. The implementation’s edge_attr tensor stores distance indices rather than chemical bond features.

ChEMBL binding and functional labels are derived from activity comments or pChEMBL: values ≥5.5 are active, values ≤4.5 inactive, and intermediate values are discarded. Contradictory molecule–assay pairs are removed, and each assay requires at least 500 molecules.

**Table A1.** Encoder and pretraining configuration.

| Quantity | Value |
| --- | --- |
| Encoder parameters | 93,484,800 |
| Layers / hidden / heads | 12 / 768 / 12 |
| Feed-forward dimension | 3,072 |
| Atom vocabulary | 211 |
| Generator layers / hidden | 3 / 256 |
| Selected-token probability | 0.25 |
| Generator replacement | argmax, unconstrained |
| RTD loss weight $\lambda$ | 50 |
| Relative attention | content-to-position |
| Absolute-position capacity | 512 |
| Relative-position embedding rows | 1,024 |
| Relative-distance bucketing | none |
| Step-1/2 heavy-atom cap | 96 |
| Position-biased input | enabled |
| Molecules > 96 heavy atoms in Step 1/2 | filtered, not truncated |
| Downstream 96-atom filter | none |
| Shared attention keys | disabled |
| Relative embedding norm | none |
| Step-1 presentations | 1,538,240,768 |
| Step-2 molecules / assays | 511,898 / 642 |
| Evaluated Step-2 snapshot | 50,000 steps |

The Step-1 exposure count is reconstructed from training manifests. Because interrupted jobs restarted with a shuffled stream, 1.538B denotes molecular presentations rather than unique molecules. Training comprised 2,500,000 updates at global batch 256 (640M presentations), 500,000 resumed updates at global batch 192 (96M presentations), and 3,133,753 exposure-correcting continuation updates at global batch 256 (802,240,768 presentations), for 6,133,753 optimizer updates in total. The final continuation ran in mixed precision on four GPUs with per-device batch 32 and gradient accumulation 2. It used AdamW with peak learning rate 5 ×10^−5^, weight decay 0.01, 10,000 warmup steps, cosine decay, and gradient-norm clipping at 1.0.

Step 2 used four-GPU distributed training with per-device batch 32, gradient accumulation 4, and global batch 512. Adam used learning rate 5 ×10^−6^, zero weight decay, 10,000 steps of linear warmup followed by linear decay, dropout 0.1, and gradient-norm clipping at 1.0. Training continued to 80,000 steps; the 50,000-step encoder was fixed for principal downstream experiments before task aggregation. The retained records identify four GPUs for the final Step-1 continuation and Step 2 but do not preserve accelerator model, wall time, GPU-hours, energy, or complete hardware accounting for earlier Step-1 phases; we report these quantities as unavailable rather than reconstructing them.

### Step-2 Label Matrix and Snapshot Sensitivity

This appendix describes the sparsity and class balance of the bioactivity labels. The principal downstream experiments use the fixed 50,000-step snapshot; no systematic downstream comparison of snapshots is reported for the final model.

The final Step-2 matrix contains 511,898 molecules and 642 assays, or 328,638,516 possible molecule–assay cells. Of these, 4,536,721 are observed (1.3805% density): 1,097,029 positive (24.18%) and 3,439,692 negative (75.82%).

### OpenADMET Data, Splits, and Evaluation

In source assay codes, HLM, MLM, and RLM mean human, mouse, and rat liver microsomes; MLM elsewhere denotes masked language modeling. The main results spell out the clearance assays to distinguish them.

Full endpoint results are reported in Table A2.

### Statistical analysis

All paired differences use GRASP minus comparator MAE. We bootstrap endpoint means with 100,000 resamples and use exact two-sided Wilcoxon and sign tests. The six baseline contrasts form one Holm family for each test; the three adaptation contrasts form another. Size and scaffold associations use endpoint averages and Holm correction over four associations per validation or test family.

The three outer assignments are stored as split1–split3. Within each outer-training partition, complete clusters are allocated to a 15% validation subset with seeds 7301, 7302, and 7303. Neural fit seeds are 9200–9202, 9300–9302, and 9400–9402 for frozen, LoRA, and full adaptation in the three splits. Training targets are standardized from inner-training rows only and predictions are inverse-transformed before MAE is computed. Table A4 reports executed row counts, target transforms, and test sizes; raw MAEs therefore remain endpoint-specific quantities and are macro-averaged only after each endpoint is scored.

### Classical baseline search

For each feature type, the random-forest grid uses 500 trees, maximum-feature choices sqrt or 0.33, and minimum leaf sizes 1, 2, or 4 (six configurations). The LightGBM grid uses 600 trees, 31 or 63 leaves, minimum child counts 10 or 20, and learning rates 0.03 or 0.06 (eight configurations), row subsampling is 0.9, feature subsampling 0.8, and L2 regularization 1.0. Validation MAE selects one configuration per endpoint and split. Chemprop 2.3.0 uses its fixed default D-MPNN architecture with a single model, no ensembling, batch size 64, a 60-epoch cap, and validation-loss early stopping with patience 12.

### Molecule size handling

The 96-heavy-atom limit was applied during pretraining but was not imposed globally on OpenADMET. Downstream molecules were neither truncated nor filtered by size: all atom tokens were retained, and variable-length inputs were padded only to the longest molecule within each batch. The encoder has an absolute-position capacity of 512 tokens. Across the 43,881 OpenADMET endpoint rows, only one row exceeded the pretraining limit: a PXR molecule with 123 heavy atoms. This molecule occurred in the inner-training partition in all three PXR splits and never in a validation or test partition. Consequently, all reported OpenADMET test sets contain molecules with at most 96 heavy atoms.

**Table A2.** Per-endpoint MAE on OpenADMET-23 (lower is better). Each entry reports the mean ± population standard deviation across three cluster-held-out splits. These deviations measure split-to-split variation, not repeated-seed variation on a fixed split. CheMeleon was initialized from its released foundation checkpoint and fully fine-tuned. **Bold** marks the best mean and <u>underlining</u> marks the second-best mean per endpoint. Rankings are based on unrounded means. Δ = 100(MAE_competitor_ MAE_GRASP_)*/*MAE_competitor_. The competitor is the runner-up when GRASP is best and the overall best method otherwise. Positive values favor GRASP; negative values favor the comparator. All methods were evaluated on identical partitions.

| Endpoint | GRASP FT | CheMeleon FT | ECFP4-LGBM | RDKit-LGBM | Chemprop | $\Delta$ (%) |
| --- | --- | --- | --- | --- | --- | --- |
| **ASAP** |  |  |  |  |  |  |
| Human liver clearance | $0.36 \pm .07$ | $0.38 \pm .04$ | **$0.33 \pm .07$** | $0.38 \pm .03$ | $0.44 \pm .06$ | -7.2 |
| Solubility | **$0.32 \pm .05$** | $0.38 \pm .07$ | <u>$0.33 \pm .07$</u> | $0.36 \pm .10$ | $0.44 \pm .09$ | +4.3 |
| LogD | **$0.47 \pm .06$** | <u>$0.55 \pm .12$</u> | <u>$0.70 \pm .19$</u> | $0.66 \pm .17$ | $0.83 \pm .24$ | +13.9 |
| MDR1 efflux | **$0.32 \pm .03$** | $0.35 \pm .06$ | <u>$0.34 \pm .04$</u> | $0.38 \pm .02$ | $0.44 \pm .04$ | +6.0 |
| MERS-CoV potency | <u>$0.56 \pm .11$</u> | $0.60 \pm .07$ | **$0.55 \pm .09$** | $0.57 \pm .10$ | $0.63 \pm .05$ | -0.5 |
| Mouse liver clearance | $0.49 \pm .05$ | <u>$0.46 \pm .03$</u> | $0.48 \pm .08$ | **$0.46 \pm .07$** | $0.51 \pm .04$ | -7.1 |
| SARS-CoV-2 potency | $0.62 \pm .05$ | <u>$0.58 \pm .06$</u> | **$0.55 \pm .02$** | $0.58 \pm .05$ | $0.64 \pm .06$ | -12.6 |
| **Biogen** |  |  |  |  |  |  |
| Human liver clearance | <u>$0.33 \pm .01$</u> | **$0.33 \pm .01$** | $0.36 \pm .01$ | $0.35 \pm .01$ | $0.34 \pm .01$ | -0.6 |
| Human plasma binding | **$0.47 \pm .13$** | $0.49 \pm .12$ | $0.48 \pm .17$ | <u>$0.48 \pm .12$</u> | $0.61 \pm .24$ | +1.7 |
| MDR1 efflux | <u>$0.34 \pm .02$</u> | **$0.33 \pm .04$** | $0.37 \pm .02$ | $0.37 \pm .01$ | $0.36 \pm .02$ | -0.8 |
| Rat liver clearance | **$0.38 \pm .02$** | <u>$0.38 \pm .03$</u> | $0.47 \pm .03$ | $0.42 \pm .03$ | $0.41 \pm .02$ | +1.5 |
| Rat plasma binding | **$0.45 \pm .06$** | <u>$0.45 \pm .04$</u> | $0.47 \pm .03$ | $0.50 \pm .01$ | $0.51 \pm .03$ | +0.5 |
| Solubility | $0.37 \pm .00$ | **$0.34 \pm .01$** | $0.38 \pm .04$ | $0.39 \pm .05$ | <u>$0.35 \pm .02$</u> | -8.8 |
| **ExpansionRx** |  |  |  |  |  |  |
| Caco-2 efflux | <u>$0.23 \pm .01$</u> | $0.24 \pm .01$ | **$0.23 \pm .00$** | $0.23 \pm .02$ | $0.25 \pm .01$ | -1.1 |
| Caco-2 permeability | **$0.29 \pm .02$** | $0.37 \pm .06$ | $0.33 \pm .03$ | <u>$0.32 \pm .03$</u> | $0.41 \pm .03$ | +9.4 |
| Human liver clearance | $0.35 \pm .02$ | **$0.35 \pm .00$** | $0.35 \pm .00$ | <u>$0.37 \pm .01$</u> | $0.37 \pm .00$ | -0.8 |
| Solubility | <u>$0.42 \pm .02$</u> | **$0.41 \pm .04$** | $0.53 \pm .10$ | $0.53 \pm .08$ | $0.43 \pm .04$ | -0.5 |
| LogD | <u>$0.27 \pm .02$</u> | **$0.26 \pm .00$** | $0.38 \pm .02$ | $0.41 \pm .03$ | $0.28 \pm .01$ | -4.1 |
| Brain binding | <u>$0.25 \pm .01$</u> | **$0.25 \pm .02$** | $0.30 \pm .02$ | $0.28 \pm .01$ | $0.30 \pm .04$ | -2.3 |
| Muscle binding | $0.26 \pm .01$ | **$0.23 \pm .02$** | <u>$0.25 \pm .04$</u> | $0.27 \pm .03$ | $0.40 \pm .02$ | -10.1 |
| Mouse liver clearance | $0.34 \pm .03$ | **$0.33 \pm .02$** | <u>$0.34 \pm .03$</u> | $0.37 \pm .01$ | $0.35 \pm .02$ | -1.9 |
| Plasma binding | **$0.24 \pm .03$** | <u>$0.25 \pm .02$</u> | $0.29 \pm .01$ | $0.30 \pm .00$ | $0.29 \pm .01$ | +0.9 |
| **PXR** |  |  |  |  |  |  |
| PXR activity | **$0.49 \pm .03$** | <u>$0.51 \pm .01$</u> | $0.55 \pm .03$ | $0.52 \pm .06$ | $0.52 \pm .03$ | +3.2 |
| **Mean** | **$0.37 \pm .00$** | <u>$0.38 \pm .01$</u> | $0.41 \pm .01$ | $0.41 \pm .01$ | $0.44 \pm .02$ | |

#### A.1.1 Overlap between ChEMBL adaptation and OpenADMET

The ChEMBL adaptation stage and OpenADMET contain a modest number of shared molecular structures. We compared the 14,261 unique standardized molecules in OpenADMET-23 against the 511,886 valid standardized molecules used during ChEMBL adaptation. Molecules were standardized using molecular cleanup, fragment-parent selection, uncharging, and sanitization.

We considered four increasingly broad definitions of exposure: (i) identical standardized structures; (ii) identical molecular connectivity, measured using the connectivity block of the InChIKey and therefore ignoring stereochemistry and protonation; and ECFP4 neighbours with Tanimoto similarity (iii) ≥ 0.90 or (iv) ≥ 0.70. These definitions provide progressively broader notions of structural exposure, but they are evaluated independently and their counts should not be added together.

**Table A3.** Paired baseline contrasts. Differences and 95% bootstrap intervals use 23 endpoints. Wilcoxon and sign-test *p*-values are Holm-adjusted within separate six-comparison families.

| Comparator | FT minus baseline | 95% CI | Wilcoxon $p$ | Sign $p$ |
| --- | --- | --- | --- | --- |
| ECFP4-LightGBM | -0.0334 | [-0.0587, -0.0116] | 0.0060 | 0.0931 |
| RDKit-LightGBM | -0.0377 | [-0.0590, -0.0192] | 0.00082 | 0.00026 |
| RDKit random forest | -0.0564 | [-0.0813, -0.0348] | 0.000063 | 0.00026 |
| Chemprop | -0.0654 | [-0.1008, -0.0375] | 0.000030 | 0.000029 |
| ECFP4 random forest | -0.0783 | [-0.1114, -0.0490] | 0.000041 | 0.00098 |
| Training median | -0.2177 | [-0.2845, -0.1612] | 0.0000014 | 0.0000014 |

**Table A4.** OpenADMET dataset and split accounting. Positive-log transforms are applied before modeling; identity denotes the retained source target. Test sizes correspond to the three stored outer cluster assignments; the remaining outer-training rows are divided into inner train and 15% cluster-held-out validation sets.

| Family | Endpoint | Rows | Target transform | Test rows (splits 1/2/3) |
| --- | --- | --- | --- | --- |
| ASAP | HLM | 299 | positive log<sub>10</sub> | 32/23/23 |
| ASAP | KSOL | 365 | positive log<sub>10</sub> | 53/48/49 |
| ASAP | LogD | 352 | identity | 55/69/59 |
| ASAP | MDR1 efflux | 424 | positive log<sub>10</sub> | 56/65/51 |
| ASAP | MERS-CoV potency | 901 | identity | 81/118/118 |
| ASAP | MLM | 300 | positive log<sub>10</sub> | 62/51/70 |
| ASAP | SARS-CoV-2 potency | 842 | identity | 69/116/50 |
| Biogen ADME | HLM CLint | 3,087 | identity | 104/75/75 |
| Biogen ADME | HPPB | 194 | identity | 25/20/21 |
| Biogen ADME | MDR1 efflux | 2,642 | identity | 77/75/76 |
| Biogen ADME | RLM CLint | 3,054 | identity | 107/124/104 |
| Biogen ADME | RPPB | 168 | identity | 20/20/20 |
| Biogen ADME | solubility | 2,173 | identity | 98/113/125 |
| ExpansionRx | Caco-2 efflux | 2,161 | positive log<sub>10</sub> | 339/260/208 |
| ExpansionRx | Caco-2 permeability | 2,156 | positive log<sub>10</sub> | 238/214/202 |
| ExpansionRx | HLM CLint | 3,595 | positive log<sub>10</sub> | 443/409/426 |
| ExpansionRx | KSOL | 5,128 | positive log<sub>10</sub> | 608/496/520 |
| ExpansionRx | LogD | 5,039 | identity | 534/570/551 |
| ExpansionRx | MBPB | 973 | positive log<sub>10</sub> | 120/113/176 |
| ExpansionRx | MGMB | 221 | positive log<sub>10</sub> | 15/22/44 |
| ExpansionRx | MLM CLint | 4,375 | positive log<sub>10</sub> | 617/528/655 |
| ExpansionRx | MPPB | 1,292 | positive log<sub>10</sub> | 133/147/146 |
| PXR | PXR activity | 4,140 | identity | 237/225/228 |

Structural matching alone does not establish whether a ChEMBL assay is semantically related to a downstream OpenADMET endpoint. During adaptation, matched molecules are associated with labels from the 642 selected ChEMBL binding or functional assays rather than with the OpenADMET records used for downstream evaluation. Nevertheless, structural exposure, and potentially related biological supervision, could make a matched molecule easier to transfer. We therefore test whether structurally exposed molecules account for the reported OpenADMET performance.

We removed exposed molecules from each test partition and recomputed MAE using the stored predictions. Models were not retrained, and the same row filters were applied to GRASP and ECFP4-LightGBM. The original GRASP mean endpoint MAE was 0.3742. Removing exact matches reduced it to 0.3724, while removing every test molecule with an ECFP4 neighbour of similarity at least 0.70 reduced it to 0.3693. Thus, performance did not deteriorate when previously seen molecules were excluded.

GRASP also retained its advantage over ECFP4-LightGBM under every exclusion rule: the filtered mean MAE difference ranged from −0.0330 to −0.0355, with GRASP winning on 16–17 of the 23 endpoints. We therefore find no evidence that the reported OpenADMET advantage is caused by direct molecule exposure during ChEMBL adaptation. This analysis tests structural exposure; it does not determine whether individual ChEMBL assays measure properties semantically related to particular OpenADMET endpoints.

**Table A5.** Executed OpenADMET downstream configuration. All three GRASP adaptation modes use a 512-unit endpoint head, AdamW, Smooth-L1 training loss, a 10% linear warmup, gradient clipping at 1.0, and validation-MAE early stopping.

| Setting | Frozen mix | LoRA mix | Full mix |
| --- | --- | --- | --- |
| Learning rate | $10^{-3}$ | $5 \times 10^{-5}$ | $10^{-5}$ |
| Epoch cap / patience |  | 60 / 12 epochs |  |
| Batch size / weight decay | | 64 / $10^{-4}$ | |
| Precision | bfloat16 when CUDA is available |  |  |
| Mixed layers |  | 4, 8, 10, 12 |  |
| LoRA rank / $\alpha$ / dropout | – | 16 / 32 / 0.1 | – |
| Trainable encoder | no | query/value LoRA | yes |

**Table A6.** Adaptation parameter counts verified across all 69 endpoint-split artifacts. LoRA adds adapter parameters, so its total parameter count differs slightly from the base encoder.

| Method | Trainable | Total | Trainable fraction |
| --- | --- | --- | --- |
| Full fine-tuning | 93,881,350 | 93,881,350 | 100.00% |
| LoRA query/value | 986,374 | 94,471,174 | 1.04% |
| Frozen layer mixing | 396,550 | 93,881,350 | 0.42% |

**Table A7.** OpenADMET adaptation by endpoint (mean MAE across three reconstructed cluster splits; lower is better). FT updates the full encoder, frozen mixing learns a readout over fixed encoder layers, and LoRA updates low-rank query/value adapters.

| Endpoint | FT | Frozen mixing | LoRA |
| --- | --- | --- | --- |
| Caco-2 permeability | $0.294 \pm 0.018$ | 0.301 | 0.297 |
| Caco-2 efflux | $0.233 \pm 0.010$ | 0.233 | 0.231 |
| ExpansionRx LogD | $0.272 \pm 0.017$ | 0.312 | 0.306 |
| ExpansionRx KSOL | $0.416 \pm 0.024$ | 0.468 | 0.415 |
| ExpansionRx HLM | $0.350 \pm 0.024$ | 0.366 | 0.369 |
| ExpansionRx MLM | $0.340 \pm 0.026$ | 0.354 | 0.364 |
| MBPB | $0.250 \pm 0.010$ | 0.253 | 0.255 |
| MGMB | $0.258 \pm 0.011$ | 0.290 | 0.290 |
| MPPB | $0.244 \pm 0.028$ | 0.265 | 0.256 |
| MERS-CoV potency | $0.557 \pm 0.109$ | 0.609 | 0.576 |
| SARS-CoV-2 potency | $0.617 \pm 0.050$ | 0.611 | 0.631 |
| ASAP LogD | $0.471 \pm 0.064$ | 0.457 | 0.454 |
| ASAP KSOL | $0.315 \pm 0.045$ | 0.331 | 0.334 |
| ASAP HLM | $0.355 \pm 0.072$ | 0.308 | 0.330 |
| ASAP MLM | $0.491 \pm 0.050$ | 0.433 | 0.477 |
| ASAP MDR1 | $0.323 \pm 0.030$ | 0.346 | 0.333 |
| PXR activity | $0.493 \pm 0.032$ | 0.508 | 0.483 |
| Biogen solubility | $0.373 \pm 0.003$ | 0.385 | 0.369 |
| Biogen HLM | $0.327 \pm 0.008$ | 0.327 | 0.333 |
| Biogen RLM | $0.376 \pm 0.024$ | 0.401 | 0.377 |
| Biogen HPPB | $0.469 \pm 0.133$ | 0.476 | 0.450 |
| Biogen RPPB | $0.445 \pm 0.060$ | 0.486 | 0.480 |
| Biogen MDR1 | $0.335 \pm 0.023$ | 0.380 | 0.329 |
| Mean, all 23 | **$0.374 \pm 0.003$** | 0.387 | 0.380 |

**Table A8.**
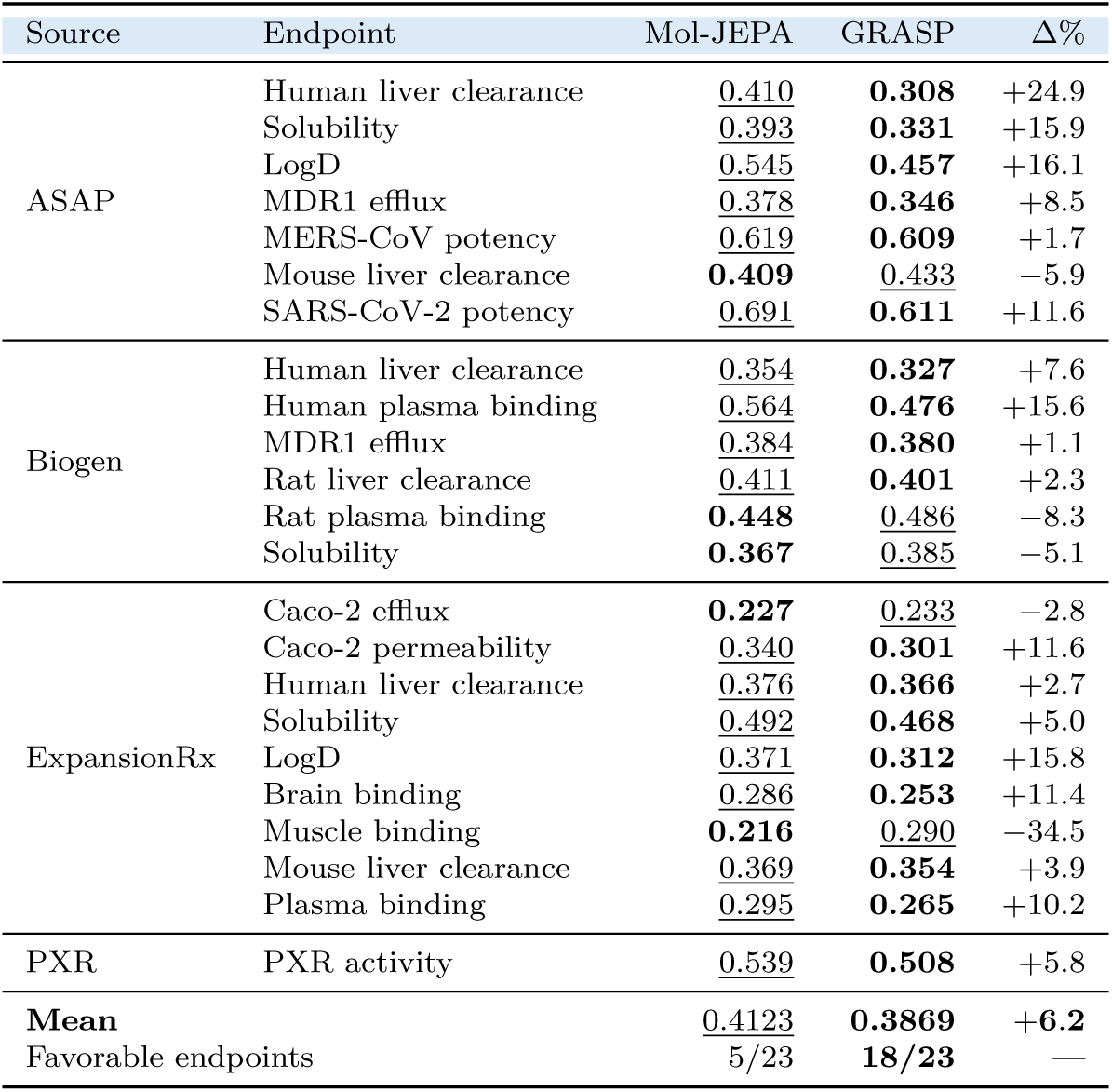
Protocol-matched frozen-encoder comparison on OpenADMET-23 (lower is better). Mol-JEPA uses its released frozen CLS representation, while GRASP uses its frozen encoder with learned layer mixing and atom-attention pooling. Both representations use the same regression MLP and the same three reconstructed cluster-held-out splits. **Bold** marks the lower MAE and <u>underlining</u> marks the other result. Δ% = 100(Mol-JEPA − GRASP)*/*Mol-JEPA; positive values favor GRASP.

**Table A9.** Structural overlap between OpenADMET-23 and the ChEMBL adaptation data. OpenADMET contains 14,261 unique standardized molecules and 43,881 labelled endpoint rows. “Test rows” pools the 12,021 test instances across the 23 endpoints and three cluster-held-out splits. A molecule can occur in more than one endpoint, so molecule and row counts differ.

| Overlap rule | Unique molecules |  | Labelled rows |  | Test rows |  |
| --- | --- | --- | --- | --- | --- | --- |
|  | Count | Fraction | Count | Fraction | Count | Fraction |
| Standardized exact | 577 | 4.05% | 1,110 | 2.53% | 203 | 1.69% |
| Same connectivity | 662 | 4.64% | 1,280 | 2.92% | 219 | 1.82% |
| ECFP4 Tanimoto $\geq 0.90$ | 674 | 4.73% | 1,301 | 2.96% | 228 | 1.90% |
| ECFP4 Tanimoto $\geq 0.70$ | 1,174 | 8.23% | 2,500 | 5.70% | 428 | 3.56% |

**Table A10.** OpenADMET performance after excluding molecules represented during ChEMBL adaptation. Δ_filter_ is filtered minus original GRASP MAE; negative values indicate lower error after exclusion. The confidence interval was obtained by bootstrapping the 23 endpoint-level differences 100,000 times. The GRASP–ECFP4 column is evaluated on the same retained rows; negative values favor GRASP.

| Exclusion rule | Endpoints exposed | Filtered MAE | $\Delta_{\text{filter}}$ | 95% CI | GRASP-ECFP4 |
| --- | --- | --- | --- | --- | --- |
| Standardized exact | 13/23 | 0.3724 | -0.0018 | [-0.0062, +0.0019] | -0.0333 |
| Same connectivity | 14/23 | 0.3721 | -0.0021 | [-0.0065, +0.0016] | -0.0336 |
| ECFP4 Tanimoto $\geq 0.90$ | 17/23 | 0.3724 | -0.0018 | [-0.0064, +0.0025] | -0.0330 |
| ECFP4 Tanimoto $\geq 0.70$ | 22/23 | 0.3693 | -0.0049 | [-0.0131, +0.0017] | -0.0355 |

**Table A11.** TDC ADMET evaluation tasks. *N* is the total number of labelled rows across the fixed train–validation pool and test set. AUROC, AUPRC, and Spearman correlation are maximized; MAE is minimized.

| Task | Type | Metric | $N$ |
| --- | --- | --- | --- |
| Ames | Classification | AUROC | 7,278 |
| BBB | Classification | AUROC | 2,030 |
| Bioavailability | Classification | AUROC | 640 |
| CYP3A4 substrate | Classification | AUROC | 670 |
| DILI | Classification | AUROC | 475 |
| HIA | Classification | AUROC | 578 |
| P-glycoprotein | Classification | AUROC | 1,218 |
| hERG | Classification | AUROC | 655 |
| CYP2C9 inhibition | Classification | AUPRC | 12,092 |
| CYP2C9 substrate | Classification | AUPRC | 669 |
| CYP2D6 inhibition | Classification | AUPRC | 13,130 |
| CYP2D6 substrate | Classification | AUPRC | 667 |
| CYP3A4 inhibition | Classification | AUPRC | 12,328 |
| Caco-2 permeability | Regression | MAE | 910 |
| LD50 | Regression | MAE | 7,385 |
| Lipophilicity | Regression | MAE | 4,200 |
| Plasma protein binding | Regression | MAE | 2,790 |
| Solubility | Regression | MAE | 9,982 |
| Hepatocyte clearance | Regression | Spearman | 1,213 |
| Microsomal clearance | Regression | Spearman | 1,102 |
| Half-life | Regression | Spearman | 667 |
| Volume of distribution | Regression | Spearman | 1,130 |

### A.2 TDC Data, Splits, and Evaluation

#### Data

We evaluate molecular-property transfer using the Therapeutics Data Commons (TDC) ADMET benchmark group [15]. The benchmark contains 22 endpoint-specific datasets spanning absorption, distribution, metabolism, excretion, and toxicity. Thirteen endpoints are binary classification tasks and nine are regression tasks. We retain the task-specific evaluation metric specified by TDC: AUROC for eight classification tasks, AUPRC for five classification tasks, MAE for five regression tasks, and Spearman correlation for four regression tasks. Table A11 lists the complete evaluation panel.

The 22 datasets contain 81,809 labelled endpoint–molecule rows in total. This is a sum over endpoints rather than a count of unique molecules because the same molecule can occur in more than one endpoint dataset. Labels are used on the scale supplied by TDC; we do not standardize regression targets or apply additional target transformations.

All SMILES in the evaluated benchmark files are RDKit-valid. The downstream loader does not apply a global heavy-atom filter and does not truncate molecules. Molecules larger than the 96-heavy-atom limit used to construct the pretraining data are processed as supplied, provided that they fit within the encoder’s positional capacity. Across the 22 task files, 70 endpoint-level rows contain more than 96 heavy atoms. Thus, the 96-heavy-atom threshold is a pretraining-data restriction rather than a downstream TDC exclusion rule.

### Splits

For every endpoint, we use the fixed test set distributed with the TDC ADMET_Group benchmark. This test membership is held constant across the final-model GRASP evaluation runs. The historical frozen-encoder comparison is described separately in section . The test set is not used for hyperparameter selection, checkpoint selection, or early stopping.

TDC provides a combined training–validation pool in addition to its fixed test set. We apply TDC’s default scaffold-splitting procedure to this pool with seeds 0–4, producing five independently seeded scaffold train–validation partitions. The nominal division of the training–validation pool is 87.5% training and 12.5% validation, corresponding to a nominal overall 70/10/20 train/validation/test allocation. Exact proportions can differ because complete molecular scaffolds are assigned together. These are five scaffold-resampled validation partitions, not five mutually exclusive folds of conventional cross-validation.

For each endpoint, learning rate and dropout are selected using only the mean validation performance across these five partitions. The search space is

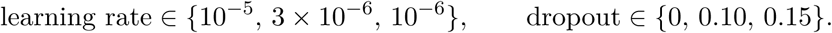

The selected configuration maximizes the endpoint’s validation metric. For MAE tasks, this is implemented by maximizing negative MAE. Consequently, each endpoint requires 3 ×3 ×5 = 45 validation runs before its final test configuration is fixed.

### Model and optimization

Molecules are represented using the same radius-0 atom-environment tokenization used by GRASP. A [CLS] token is added to each molecular sequence. The final encoder-layer representation of this token is passed through a task-specific dropout–linear–GELU–dropout–linear prediction head. Unless a result is explicitly identified as a frozen-encoder experiment, the complete encoder and prediction head are fine-tuned jointly.

Classification models are optimized with binary cross-entropy on the logits. Regression models are optimized with mean squared error, irrespective of whether their reporting metric is MAE or Spearman correlation. We use Adam with zero weight decay, a batch size of 32, and gradient-norm clipping at 1.0. Training runs for at most 100 epochs. The learning rate increases linearly during the first 10% of optimization steps and remains constant after warmup. Validation is performed every five epochs, and the checkpoint with the best task-specific validation metric is retained.

### Locked-test evaluation

After the learning rate and dropout have been fixed, we run three independent downstream training runs with seeds 100, 101, and 102. These runs use scaffold train–validation partitions generated with split seeds 0, 1, and 2, respectively, while preserving exactly the same fixed test membership. The test set is evaluated only after the best validation checkpoint for a run has been selected.

For each endpoint, we report the arithmetic mean and population standard deviation across the three test runs,

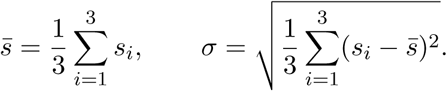

The reported deviation therefore reflects both downstream-training randomness and variation among the three scaffold train–validation assignments; it is not pretraining-seed uncertainty.

AUROC, AUPRC, and Spearman correlation are reported with higher values indicating better performance, whereas lower MAE is better. Because the endpoints use different metrics and, for MAE, different target scales, we do not compute a numerical average across the 22 TDC tasks. Comparisons are instead summarized using individual task results and the number of tasks on which each method has the favorable point estimate. ClinTox is not part of this 22-task evaluation panel and is excluded from its task counts.

**Table A12.** RTD versus MLM on TDC in the controlled 415M-molecule configuration. Values are mean ± population standard deviation over three seeds. **Bold** marks the favorable point estimate. BBB is excluded because a valid MLM result was not available.

| Task | Metric | RTD-15%-S2 | MLM-15%-S2 |
| --- | --- | --- | --- |
| **Absorption** |  |  |  |
| Caco-2 | MAE ↓ | **0.35 ± .02** | 0.36 ± .03 |
| Bioavailability | AUROC ↑ | **0.68 ± .02** | 0.63 ± .04 |
| Lipophilicity | MAE ↓ | **0.46 ± .01** | 0.51 ± .01 |
| Solubility | MAE ↓ | **0.80 ± .02** | 0.81 ± .01 |
| HIA | AUROC ↑ | **0.97 ± .01** | 0.96 ± .02 |
| P-glycoprotein | AUROC ↑ | 0.92 ± .01 | **0.93 ± .01** |
| **Distribution** |  |  |  |
| BBB Martins | AUROC ↑ | **0.88 ± .02** | 0.86 ± .02 |
| PPBR | MAE ↓ | **7.17 ± .28** | 7.49 ± .15 |
| VDss | Spearman ↑ | **0.56 ± .10** | 0.50 ± .14 |
| **Metabolism** |  |  |  |
| CYP2C9 inhibition | AUPRC ↑ | **0.77 ± .02** | 0.74 ± .03 |
| CYP2D6 inhibition | AUPRC ↑ | **0.69 ± .01** | 0.63 ± .02 |
| CYP3A4 inhibition | AUPRC ↑ | **0.87 ± .01** | 0.86 ± .01 |
| CYP2C9 substrate | AUPRC ↑ | 0.32 ± .02 | **0.40 ± .03** |
| CYP2D6 substrate | AUPRC ↑ | 0.54 ± .04 | **0.57 ± .04** |
| CYP3A4 substrate | AUROC ↑ | 0.65 ± .01 | **0.67 ± .03** |
| **Excretion** |  |  |  |
| Half-life | Spearman ↑ | **0.29 ± .09** | 0.19 ± .22 |
| Hepatocyte clearance | Spearman ↑ | **0.43 ± .02** | 0.38 ± .05 |
| Microsomal clearance | Spearman ↑ | **0.59 ± .02** | 0.47 ± .06 |
| **Toxicity** |  |  |  |
| LD50 | MAE ↓ | **0.54 ± .01** | 0.55 ± .02 |
| hERG | AUROC ↑ | **0.82 ± .02** | 0.75 ± .01 |
| Ames | AUROC ↑ | **0.80 ± .01** | 0.79 ± .01 |
| DILI | AUROC ↑ | 0.87 ± .01 | **0.88 ± .01** |
| **Favorable tasks** | **Count** | **17/22** | 5/22 |

**Table A13.** Effect of ChEMBL adaptation on the 22-task TDC ADMET panel. GRASP-S1 uses structural pretraining only; GRASP also includes ChEMBL adaptation. Values are mean ± population standard deviation over three downstream seeds following validation-only hyperparameter selection. Bold marks the favorable unrounded mean within each task. BBB is included and ClinTox is excluded. GRASP has the favorable point estimate on 19 tasks and GRASP-S1 on three.

| Task | Metric | GRASP-S1 | GRASP |
| --- | --- | --- | --- |
| Caco-2 | MAE ↓ | 0.470 ± 0.040 | **0.389 ± 0.009** |
| Bioavailability | AUROC ↑ | 0.649 ± 0.046 | **0.687 ± 0.022** |
| Lipophilicity | MAE ↓ | 0.449 ± 0.011 | **0.406 ± 0.003** |
| Solubility | MAE ↓ | 0.794 ± 0.003 | **0.736 ± 0.009** |
| HIA | AUROC ↑ | **0.960 ± 0.004** | 0.953 ± 0.017 |
| P-glycoprotein | AUROC ↑ | 0.915 ± 0.011 | **0.919 ± 0.007** |
| BBB | AUROC ↑ | 0.874 ± 0.003 | **0.880 ± 0.022** |
| PPBR | MAE ↓ | 8.361 ± 0.394 | **7.300 ± 0.346** |
| VDss | Spearman ↑ | 0.599 ± 0.025 | **0.665 ± 0.019** |
| CYP2C9 inhibition | AUPRC ↑ | 0.774 ± 0.004 | **0.791 ± 0.003** |
| CYP2D6 inhibition | AUPRC ↑ | 0.624 ± 0.009 | **0.716 ± 0.017** |
| CYP3A4 inhibition | AUPRC ↑ | 0.853 ± 0.011 | **0.891 ± 0.005** |
| CYP2C9 substrate | AUPRC ↑ | 0.385 ± 0.014 | **0.405 ± 0.026** |
| CYP2D6 substrate | AUPRC ↑ | 0.636 ± 0.024 | **0.640 ± 0.034** |
| CYP3A4 substrate | AUROC ↑ | 0.565 ± 0.046 | **0.674 ± 0.035** |
| Half-life | Spearman ↑ | **0.509 ± 0.055** | 0.455 ± 0.062 |
| Hepatocyte clearance | Spearman ↑ | 0.408 ± 0.042 | **0.471 ± 0.018** |
| Microsomal clearance | Spearman ↑ | 0.556 ± 0.048 | **0.602 ± 0.017** |
| LD50 | MAE ↓ | 0.576 ± 0.002 | **0.486 ± 0.002** |
| hERG | AUROC ↑ | **0.797 ± 0.023** | 0.773 ± 0.025 |
| Ames | AUROC ↑ | 0.823 ± 0.008 | **0.833 ± 0.005** |
| DILI | AUROC ↑ | 0.846 ± 0.033 | **0.926 ± 0.021** |
| **Favorable tasks** | Count | **3/22** | **19/22** |

## References

[1] Satya Pratik Srivastava, Rohan Gorantla, Sharath Krishna Chundru, Claire JR Winkelman, Antonia SJS Mey, and Rajeev Kumar Singh. Explainable active learning framework for ligand binding affinity prediction. Digital Discovery, 5(2):769–779, 2026.

[2] Naifeng Wen, Guanqun Liu, Jie Zhang, Rubo Zhang, Yating Fu, and Xu Han. A fingerprints based molecular property prediction method using the bert model. Journal of Cheminformatics, 14(1):71, 2022.

[3] John J. Irwin, Khanh G. Tang, Jennifer Young, Chinzorig Dandarchuluun, Benjamin R. Wong, Munkhzul Khurelbaatar, Yurii S. Moroz, John Mayfield, and Roger A. Sayle. ZINC20: A free ultralarge-scale chemical database for ligand discovery. Journal of Chemical Information and Modeling, 60(12):6065–6073, 2020. doi: 10.1021/acs.jcim.0c00675. URL 10.1021/acs.jcim.0c00675.

[4] Rohan Gorantla. Machine learning in drug discovery: advancing protein-ligand binding affinity predictions. PhD thesis, 2025.

[5] Barbara Zdrazil, Eloy Felix, Fiona Hunter, Emma J. Manners, James Blackshaw, Steve Corbett, Marleen de Veij, Haralambos Ioannidis, David M. Lopez, Juan F. Mosquera, et al. The ChEMBL database in 2023: A drug discovery platform spanning multiple bioactivity data types and time periods. Nucleic Acids Research, 52(D1):D1180–D1192, 2024. doi: 10.1093/nar/gkad1004. URL 10.1093/nar/gkad1004.

[6] ChEMBL Team. ChEMBL database release 36, 2025. URL 10.6019/CHEMBL.database.36.

[7] Kevin Clark, Minh-Thang Luong, Quoc V. Le, and Christopher D. Manning. ELECTRA: Pre-training text encoders as discriminators rather than generators. In International Conference on Learning Representations, 2020. URL https://openreview.net/forum?id=r1xMH1BtvB.

[8] David Rogers and Mathew Hahn. Extended-connectivity fingerprints. Journal of Chemical Information and Modeling, 50(5):742–754, 2010. doi: 10.1021/ci100050t. URL doi.org/10.1021/ci100050t.

[9] Chengxuan Ying, Tianle Cai, Shengjie Luo, Shuxin Zheng, Guolin Ke, Di He, Yanming Shen, and Tie-Yan Liu. Do transformers really perform bad for graph representation? In Advances in Neural Information Processing Systems, volume 34, 2021. URL https://proceedings.neurips.cc/paper/2021/hash/f1c1592588411002af340cbaedd6fc33.html.

[10] Wonpyo Park, Woong-Gi Chang, Donggeon Lee, Juntae Kim, and Seung-won Hwang. GRPE: Relative positional encoding for graph transformer. In ICLR 2022 Workshop on Machine Learning for Drug Discovery, 2022.

[11] Rohan Gorantla, Aryo Pradipta Gema, Ian Xi Yang, Álvaro Serrano-Morrás, Benjamin Suutari, Jordi Juárez-Jiménez, and Antonia SJS Mey. Learning binding affinities via fine-tuning of protein and ligand language models. Journal of Chemical Information and Modeling, 65(22): 12279–12291, 2025.

[12] Hugo MacDermott-Opeskin, Jenke Scheen, Cas Wognum, Joshua T. Horton, Devany West, et al. A computational community blind challenge on pan-coronavirus drug discovery data. Journal of Chemical Information and Modeling, 66(6):3129–3149, 2026. doi: 10.1021/acs.jcim.5c02106. URL 10.1021/acs.jcim.5c02106.

[13] Maria A. Castellanos, Hugo MacDermott-Opeskin, Jon Ainsley, Jonathan A. Swain, Sean Colby, Devany West, Georgia Channing, Andrew Good, Elyse Bourque, Lakshminarayana Vogeti, Renato Skerlj, Tiansheng Wang, Mark Ledeboer, David Alencar Araripe, Rafal Bachorz, Amitesh Badkul, et al. Assessing the state of the art in admet prediction: Lessons from the openadmet-expansionrx community blind challenge. ChemRxiv, 2026(0923), 2026. doi: 10. 26434/chemrxiv.15009342/v1. URL https://chemrxiv.org/doi/abs/10.26434/chemrxiv.15009342/v1.

[14] Cheng Fang, Ye Wang, Richard Grater, Sudarshan Kapadnis, Cheryl Black, Patrick Trapa, and Simone Sciabola. Prospective validation of machine learning algorithms for absorption, distribution, metabolism, and excretion prediction: An industrial perspective. Journal of Chemical Information and Modeling, 63(11):3263–3274, 2023.

[15] Kexin Huang, Tianfan Fu, Wenhao Gao, Yue Zhao, Yusuf Roohani, Jure Leskovec, Connor W. Coley, Cao Xiao, Jimeng Sun, and Marinka Zitnik. Therapeutics data commons: Machine learning datasets and tasks for drug discovery and development. In *Proceedings of the Neural Information Processing Systems Track on Datasets and Benchmarks*, volume 1, 2021. URL https://datasets-benchmarks-proceedings.neurips.cc/paper_files/paper/2021/hash/4c56ff4ce4aaf9573aa5dff913df997a-Abstract-round1.html.

[16] Ruoxi Sun, Hanjun Dai, and Adams Wei Yu. Does gnn pretraining help molecular representation? Advances in Neural Information Processing Systems, 35:12096–12109, 2022.

[17] Maciej Sypetkowski, Frederik Wenkel, Farimah Poursafaei, Nia Dickson, Karush Suri, Philip Fradkin, and Dominique Beaini. On the scalability of gnns for molecular graphs. Advances in Neural Information Processing Systems, 37:19870–19906, 2024.

[18] Harshit Singh, Aastha Malhotra, Satya Pratik Srivastava, RAJEEV KUMAR Singh, and Rohan Gorantla. Antibody-antigen affinity prediction with chain-aware protein language modeling. bioRxiv, pages 2026–06, 2026.

[19] Yu Rong, Yatao Bian, Tingyang Xu, Weiyang Xie, Ying Wei, Wenbing Huang, and Junzhou Huang. Self-supervised graph transformer on large-scale molecular data. Advances in neural information processing systems, 33:12559–12571, 2020.

[20] Yuyang Wang, Jianren Wang, Zhonglin Cao, and Amir Barati Farimani. Molecular contrastive learning of representations via graph neural networks. Nature Machine Intelligence, 4(3): 279–287, 2022.

[21] Seyone Chithrananda, Gabriel Grand, and Bharath Ramsundar. ChemBERTa: Large-scale self-supervised pretraining for molecular property prediction, 2020. URL https://arxiv.org/abs/2010.09885.

[22] Jerret Ross, Brian Belgodere, Vijil Chenthamarakshan, Inkit Padhi, Youssef Mroueh, and Payel Das. Large-scale chemical language representations capture molecular structure and properties. Nature Machine Intelligence, 4(12):1256–1264, 2022. doi: 10.1038/s42256-022-00580-7. URL 10.1038/s42256-022-00580-7.

[23] Gengmo Zhou, Zhifeng Gao, Qiankun Ding, Hang Zheng, Hongteng Xu, Zhewei Wei, Linfeng Zhang, and Guolin Ke. Uni-Mol: A universal 3d molecular representation learning framework. In International Conference on Learning Representations, 2023. URL https://openreview.net/forum?id=6K2RM6wVqKu.

[24] Xiaohong Ji, Zhen Wang, Zhifeng Gao, Hang Zheng, Linfeng Zhang, Guolin Ke, and Weinan E. Uni-Mol2: Exploring molecular pretraining model at scale. In Advances in Neural Information Processing Systems, volume 37, 2024. URL https://papers.neurips.cc/paper_files/paper/2024/hash/53923bb44655a7defb31c7744c01b62b-Abstract-Conference.html.

[25] Jackson Burns, Akshat Zalte, and William H. Green. Descriptor-based foundation models for molecular property prediction, 2025. URL https://arxiv.org/abs/2506.15792. Preprint.

[26] Oscar Méndez-Lucio, Christos A. Nicolaou, and Berton Earnshaw. MolE: A foundation model for molecular graphs using disentangled attention. Nature Communications, 15:9431, 2024. doi: 10.1038/s41467-024-53751-y. URL 10.1038/s41467-024-53751-y.

[27] Pengcheng He, Jianfeng Gao, and Weizhu Chen. DeBERTaV3: Improving DeBERTa using ELECTRA-style pre-training with gradient-disentangled embedding sharing. In International Conference on Learning Representations, 2023. URL https://openreview.net/forum?id=sE7-XhLxHA.

[28] Jun Xia, Shaorong Chen, Yue Liu, Zhangyang Gao, Jiangbin Zheng, Xihong Yang, and Stan Z Li. Discognn: A sample-efficient framework for self-supervised graph representation learning. In 2024 IEEE 40th International Conference on Data Engineering (ICDE), pages 2876–2888. IEEE, 2024.

[29] Greg Landrum et al. RDKit: Open-source cheminformatics software, 2025. URL https://www.rdkit.org.

[30] Ashish Vaswani, Noam Shazeer, Niki Parmar, Jakob Uszkoreit, Llion Jones, Aidan N. Gomez, Lukasz Kaiser, and Illia Polosukhin. Attention is all you need. In Advances in Neural Information Processing Systems, 2017.

[31] Edward J. Hu, Yelong Shen, Phillip Wallis, Zeyuan Allen-Zhu, Yuanzhi Li, Shean Wang, Lu Wang, and Weizhu Chen. LoRA: Low-rank adaptation of large language models. In International Conference on Learning Representations, 2022. URL https://openreview.net/forum?id=nZeVKeeFYf9.

[32] Rohan Gorantla, Alzbeta Kubincova, Benjamin Suutari, Benjamin P Cossins, and Antonia SJS Mey. Benchmarking active learning protocols for ligand-binding affinity prediction. Journal of Chemical Information and Modeling, 64(6):1955–1965, 2024.

